# Tissue-wide metabolomics reveals wide impact of gut microbiota on mice metabolite composition

**DOI:** 10.1101/2021.08.12.456100

**Authors:** Iman Zarei, Ville M. Koistinen, Marietta Kokla, Anton Klåvus, Ambrin Farizah Babu, Marko Lehtonen, Seppo Auriola, Kati Hanhineva

## Abstract

The essential role of gut microbiota in health and disease is well-recognized, but the biochemical details underling beneficial impact remain largely undefined. Dysbiosis of gut bacteria results in the alteration of certain microbial and host metabolites, and identifying these markers could enhance the early detection of certain diseases. We report LC-MS based non-targeted metabolic profiling to demonstrate a large effect of gut microbiota on mammalian tissue metabolites. It was hypothesized that gut microbiota influences the overall biochemistry of the host metabolome and this effect is tissue-specific. Thirteen different tissues from germ-free and conventional mice were selected and their metabolic differences were analyzed. Our study demonstrated a large effect of the microbiome on mammalian biochemistry at different tissue levels and resulted in significant modulation of metabolites from multiple metabolic pathway (p ≤ 0.05). A vast metabolic response of host to metabolites generated by the microbiota was observed, Hundreds of molecular features were detected exclusively in one mouse group, with the majority of these being unique to specific tissue, suggesting direct impact gut microbiota on host metabolism.

## 1. Introduction

Microbiota (microbial community) or microbiome (collective genome of microbial community) is defined as the commensal, symbiotic, and pathogenic microbial community including bacteria, fungi, archaea, algae, and small protists which reside inside and on the host body [2–4]. Human microbiota comprises trillions of microorganisms and can encode significantly more individual genes than the human genome [5–7]. Some of our tissues, such as those with a mucosal membrane, contain highly adapted and evolved microbial consortia [1], with the vast majority of the microbiota within our gastrointestinal (GI) tract because of its nutrient-rich environment. Gut microbiota has a complex influence on human physiology and nutritional status [8, 9] by influencing the absorption, metabolism, and storage of ingested nutrients and by producing a diverse array of metabolites. Examples include digestion and bioconversion of food components such as hydrolysis and fermentation of indigestible plant nutrients (e.g., oligo- and polysaccharides known as microbiota-accessible carbohydrate or in short MAC) to make them bioavailable to the host [10–12] and subsequently, production of metabolites involved in energy homeostasis, namely short-chain fatty acids (SCFAs) [13, 14]; biosynthesis of indoles, aromatic amino acid metabolites, vitamins, and sphingolipids [15–17]; cholesterol synthesis inhibition and bile acid biotransformation by regulating their composition, abundance, and signaling [18–20]; stimulation and regulation of the immune system as well as inhibition of pathogens (e.g., production of antimicrobial compounds, regulating of intestinal pH, and competition for ecological niche) which in the end leads to support of intestinal function [21–23]; removal of toxins, drug residues and carcinogens from the body [24, 25]; and even potential regulation of host central nervous system [26–28].

Despite the fact that the mature microbiota is very resilient, high inter-individual variability in the composition of human gut microbiota can be explained by internal and external stimulants such as age, genotype, mode of delivery, antibiotic use, diet, demography, lifestyle, social interactions, stress, and environmental exposure to various xenobiotics [29–31]. Among the mentioned factors, diet alone is one of the most important modifiable lifestyle factors contributing to variation in gut microbiota composition, and indeed, the impact of diet on gut microbiota is found to be higher as compared to e.g., genotype [32]. Furthermore, the inter-individual variation might explain why the impact of nutritional interventions varies among individuals, even though the same food was consumed [33]. Given that the differences in commensal microbiota and even reduced microbial diversity may impact human health and disease, changes in the composition of gut microbiota are linked to the development of many disorders such as type 2 diabetes, cardiovascular dyslipidemia, and cirrhosis, cancer, allergies, inflammatory bowel disease (IBD), neurodevelopmental disorders (e.g., autism), aging, and many more [34–36]. Therefore, manipulation of gut microbiota in preventing and treating chronic diseases can lead us to a deeper systematic understanding of the microbiota-host interface, as well as take us to a closer step towards customizing dietary schemes for personalized dietary treatment by expanding to a nutrient-microbiota-host approach [37].

It has been proven difficult to establish microbe-related biomarkers for health and disease due to the current lack of knowledge on the impact of diet and other environmental factors on microbiota and its variation and function across different populations [38]. To understand this massively complex factor in human health, there is a need to model it effectively. The study of microbiota-host interactions is challenging because of the high degree of crosstalk between these two domains. Nowadays, next-generation-sequencing platforms are used to annotate bacterial species associated with the gut in higher organisms. However, profiling of complex microbial communities via 16S sequencing lacks the information on the fingerprint of the microbiome functional status and the actual activities of the microbes mediated via the metabolites they produce [39]. The identity and function of gut microbiota have a direct impact on the metabolites that are produced, many of which are still structurally uncharacterized. Many of these metabolites are taken up into host circulation and eventually into various tissues to participate in endogenous metabolism. Notably, the effect of microbial compounds within the mammalian host environment varies from one tissue to another based on the type and metabolic status of affected tissues [40–42]. Metabolomics is a well-established and powerful tool that can be applied to identify microbiome-derived or microbiome-modified metabolites and to better understand the modulation of microbiota and how it affects the metabolism in a host [43, 44]. Metabolomics can help to define the metabolic interactions among the host, diet, and gut microbiota [45].

We herein hypothesized that gut microbiota influences the overall biochemistry of the host metabolome and its effect is tissue-specific (variable for each tissue). Thus, the aim of the study was to establish a comparative metabolite-level overview of 13 different tissues from germ-free [8] and conventional mice (murine-pathogen-free, MPF) affected by the intestinal microbial community using non-targeted metabolite profiling approach. We chose to analyze multiple tissues because it provides an excellent opportunity to assess the extent of the interplay between bacterial metabolic and systemic human pathways. The metabolite composition of plasma, heart, liver, pancreas, muscle, duodenum, jejunum, ileum, cecum, colon, visceral adipose tissue (VAT), subcutaneous adipose tissue (SAT), and brown adipose tissue (BAT) were analyzed, and demonstrated a massive metabolic impact across all the tissues studied. We observed significantly large number of chemical species in the tissues because of the presence of the microbiota, and a range of 29-74% of all detectable metabolites varied in concentration by at least 50% between the 2 mouse lines.

## 2. Materials and Methods

### 2.1. Tissue sample collection and preparation

Blood, heart, liver, pancreas, muscle, duodenum, jejunum, ileum, cecum, colon, VAT, SAT, and BAT tissues from five GF and five MPF male C57BL/6NTac mice age 10 weeks were obtained from Taconic Biosciences (www.taconic.com/mouse-model/black-6-b6ntac). The sterile natural ingredient NIH #31M Rodent Diet (www.taconic.com/quality/animal-diet) was used as the standard diet. To assure the germ-free status of the mice used in the study, trimethylamine N-oxide (TMAO), a metabolite with microbial origin [46], was used as a reference compound. Blood was collected (K2-EDTA Microtainer Tubes) and centrifuged at 3,000 *g* for 10 min at room temperature. Other tissues were rinsed with phosphate-buffered saline thoroughly. Furthermore, all the tissues were snap-frozen in liquid nitrogen and shipped on dry ice. Upon arrival, samples were immediately transferred and kept at −80 °C until further processing for metabolomics.

All frozen samples were cryo-ground and then 100 ±2 mg of powdered sample was cryo-weighted into 1.5 mL Eppendorf tubes. The tissues (except plasma) were treated with 80% ice-cold methanol in a ratio of 300 μL solvent per 100 mg tissue. Then the samples were briefly vortexed and incubated on a shaker (Heidolph Multi Reax) at 2,000 rpm for 15 min at room temperature. For plasma samples, acetonitrile (ACN) was used as a solvent with the ratio of 1:4 vol:vol (plasma to solvent) and vortexed. All the samples were centrifuged for 10 min at 4°C (18,000 *g*), and the supernatant fractions were filtered using 0.2-μm Acrodisc® Syringe Filters with a PTFE membrane (PALL Corporation) and stored at −20°C until further analysis with LC-MS. The order of the samples was randomized before the analysis.

### 2.2. Instrumentation

We used ultra-high-performance liquid chromatography (1290 LC system, Agilent Technologies, Santa Clara, CA) with high-resolution mass spectrometry (6540 Q-TOF-MS, Agilent Technologies, Santa Clara, CA) for non-targeted metabolite profiling [47–49]. We used two chromatographic separation techniques, which were hydrophilic interaction chromatography (HILIC) and reversed-phase (RP), and further acquired data in both positive and negative electrospray ionization (ESI) modes, optimized for as wide metabolite coverage as possible. Aliquots of 2 μL from all the specific sample matrices were generated as a pooled quality control sample (QC) and were injected in the beginning of the analysis as well as between sample types (every 10^th^ injection). For RP analysis, the mobile phase flow rate was 500 μl/min with Zorbax RRHD Eclipse XDB-C18 column (100 × 2.1 mm, 1.8 μm; Agilent Technologies). The column temperature was maintained at 50°C. Mobile phase was 0.1% v/v formic acid in water (A) and 0.1% (v/v) formic acid in methanol (B). We used gradient elution which was as follows: 0–10 min: 2% → 100% solution B; 10–14.5 min: 100% solution B; 14.5–14.51 min: 100% → 2% solution B; and 14.51–16.5 min: 2% solution B. For HILIC analysis, mobile phase flow rate was 600 μL/min with Acquity UPLC® BEH Amide column (100 mm × 2.1 mm, 1.7 μm; Waters Corporation, Milford, MA). The column temperature was maintained at 45°C. Mobile phase was 50% v:v acetonitrile (A) and 90% v:v acetonitrile (B). Both solvents contained 20 mmol/L ammonium formate, pH 3. The following gradient elution was used: 0–2.5 min, 100% B; 2.5–10 min, 100% B→0% B; 10–10.1 min, 0% B→100% B; 10.1–14 min, 100% B. The sample injection volume was 3 μL and the sample tray temperature was kept at 4°C during the analysis.

The mass spectrometry (MS) conditions were: drying gas temperature of 325°C with a flow of 10 L/min, a sheath gas temperature of 350°C and a flow of 11 L/min, a nebulizer pressure of 45 psi (310 kPa), capillary voltage of 3,500 V, nozzle voltage of 1,000 V, fragmentor voltage of 100 V, and a skimmer voltage of 45 V. Data acquisition was performed using extended dynamic range mode (2 GHz), and the instrument was set to acquire ions over the mass range *m/z* 50–1,600. Data were collected in the centroid mode at an acquisition rate of 2.5 spectra/s (*i.e.*, 400 ms/spectrum) with an abundance threshold of 150. For automatic data-dependent MS/MS analyses, the precursor isolation width was 1.3 Da, and from every precursor scan cycle, the 4 ions with the highest abundance were selected for fragmentation with the collision energies of 10, 20, and 40V. These ions were excluded after 2 product ion spectra and released again for fragmentation after a 0.25 min hold. The precursor scan time was based on ion intensity, ending at 20,000 counts or after 300 ms. The product ion scan time was 300 ms. Continuous mass axis calibration was applied throughout the analysis using two reference ions *m/z* 121.050873 and *m/z* 922.009798 in the positive mode and *m/z* 112.985587 and *m/z* 966.000725 in the negative mode.

### 2.3. Data extraction and compound identification

Raw data was processed through MS-DIAL software (version 3.00) for baseline filtering, baseline calibration, peak picking, identification, peak alignment, and peak height integration [50]. Centroid spectra peaks higher than 400 counts were restricted to ion species [M-H]^−^ and [M+Cl]^−^ in negative and [M+H]^+^ and [M+Na]^+^ in positive modes. The mass tolerance for compound mass was ±15 mDa, retention time ±0.2 min, and symmetric expansion value ±10 mDa for chromatograms. Compounds were identified by comparison to library entries of purified standards and compared against METLIN (https://metlin.scripps.edu), MassBank of North America (MoNA, https://mona.fiehnlab.ucdavis.edu), Human Metabolome Database (HMDB, www.hmdb.ca), and LIPID MAPS (www.lipidmaps.org) metabolomics databases. The MS/MS fragmentation of the metabolites was compared with candidate molecules found in databases and verified with earlier literature on the same or similar compounds. Metabolomics Center of Biocenter Kuopio maintains an in-house library of over 600 authenticated standards that contains the retention time, mass to charge ratio (*m/z*), and chromatographic data (including MS/MS spectral data) on all molecules present in the library.

### 2.4. Statistical analysis

The combined data matrix, *i.e.*, HILIC (positive and negative ionization modes) and RP (positive and negative ionization modes) comprised 24,294 molecular features from 13 tissues from the GF and MPF mice, which underwent statistical analysis. There was a total of 5,961 and 3,946 molecular features obtained from HILIC, and 9,256 and 5,131 features from RP, in positive and negative ionization modes, respectively. Before performing any statistical analysis, the false zero values were imputed for each mouse group in a tissue, individually. An arbitrary raw abundance value of 10,000 was set as the threshold. Following rules were applied based on the signals of biological replicate measurements: (1) if the metabolite raw abundance was zero in more than 60% of the replicates, then the zero values were considered true zero. Therefore, all the non-zero values were replaced with zero regardless if they were smaller or higher than the arbitrary threshold (*i.e.*, default intensity of 10,000); (2) if the metabolite raw abundance was zero in less than 60% of the replicates and the non-zero values were lower than the threshold, then zero values were also considered true zero and the non-zero values were replaced with zero; (3) if the metabolite raw abundance was zero in less than 60% of the replicates and the non-zero values were higher than the threshold, then zero values were considered false and were replaced with an imputed value. The imputed value was calculated for each molecular feature as the average of the non-zero raw abundances in that tissue-specific mouse group.

A fold change (FC) value was calculated for each molecular feature and tissue by dividing the average signal abundance of the GF samples (“treatment” group) with that of the MPF samples (“control” group). Thus, FC > 1 signifies a higher abundance of the molecular feature in the GF mice and FC < 1 a higher abundance in the MPF mice.

Data processing was carried out by R package (3.5.3) for unscaled data. Mann–Whitney *U-*test was chosen to identify the most differentially abundant molecular features between the GF and MPF for the tissue-specific metabolite levels. False discovery rate (FDR, corrected *p*-value, *q*-value) was performed based on Benjamini and Hochberg’s method. Significant metabolites had *p*-values of ≤ 0.05 and *q*-values below the threshold of ≤ 0.05.

The raw abundances of metabolites were first z-normalized based on the following formula: 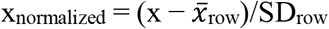. The *k*-means cluster analysis and hierarchical clustering were performed by the open-source software Multi experiment Viewer (MeV, http://mev.tm4.org) to visualize the common trends in the profile of the different molecular features (*k*-means clustering) as well as visualizing the abundance of the features when compared against the other molecular features present in the same cluster. The number of clusters was set to 10. The number of clusters was optimized based on visual inspection to reveal as many clusters as possible with a distinct pattern of molecular features without having two similar clusters.

After the statistical analysis, further filtering was applied on the obtained differential metabolic features based on the following inclusion criteria. The inclusion criteria must have existed in at least one tissue and one mouse group (*i.e.*, GF or MPF): (1) fold change ≥ 1.3 with a *p*-value ≤ 0.05, and *q*-value ≤ 0.05 (2) high-intensity metabolite values (raw abundance ≥ 100,000), (3) containment of MS/MS fragmentation, (4) and retention time ≥ 0.7 min. This filtering procedure resulted in a set of 4,605 statistically significant molecular features, which then underwent *k*-means clustering analysis. Principal component analysis (PCA) was applied to all the 24,294 molecular features for visualization of the overall metabolite feature pattern of different tissues. Tissue-wise volcano plot visualization was applied on the statistically significant molecular features to display discriminatory molecular features within a tissue between the GF and MPF mice using MetaboAnalyst platform (https://www.metaboanalyst.ca/) [51].

## 3. Results

### 3.1. The impact of microbiota on the metabolome of GF and MPF mice was tissue-wide

A total of 130 samples from 13 tissues were analyzed from five GF and five MPF mice using high-resolution LC-MS platform with four analytical modes employing RP and HILIC modes with both positive and negative ionization providing an initial wide-scale assessment of the effect of the microbiome on mammalian metabolism. Figure 1 and Supplementary Table 1 summarize the detection rate and the proportion of differential and unique molecular features in the 13 different tissues; Most of detected individual features were present in the GI tract tissues (*i.e.*, duodenum, jejunum, ileum, cecum, and colon), and the liver (≃41-74%), with the cecum containing highest percentage of all detected features, whereas plasma with 23% of all detected features showed the lowest number among all the 13 tissues. The GI tract tissues, and the liver also showed the highest percentage of significant molecular features (≃26-64%) (according to the statistical criteria applied) with the cecum containing highest percentage of significant molecular feature. It is noteworthy that the GF mice showed the highest percentage of significant molecular features (≥ 50%) in all the tissues from the GI as well as plasma and BAT when compared to the MPF mice in the same tissue. Contrarily, MPF mice showed the highest percentage of significant molecular features in the liver, SAT, pancreas, muscle, VAT, and heart. Colon and SAT were the only tissues with more than half of their detected molecular features unique to the GF and MPF, respectively.

**Figure 1.**
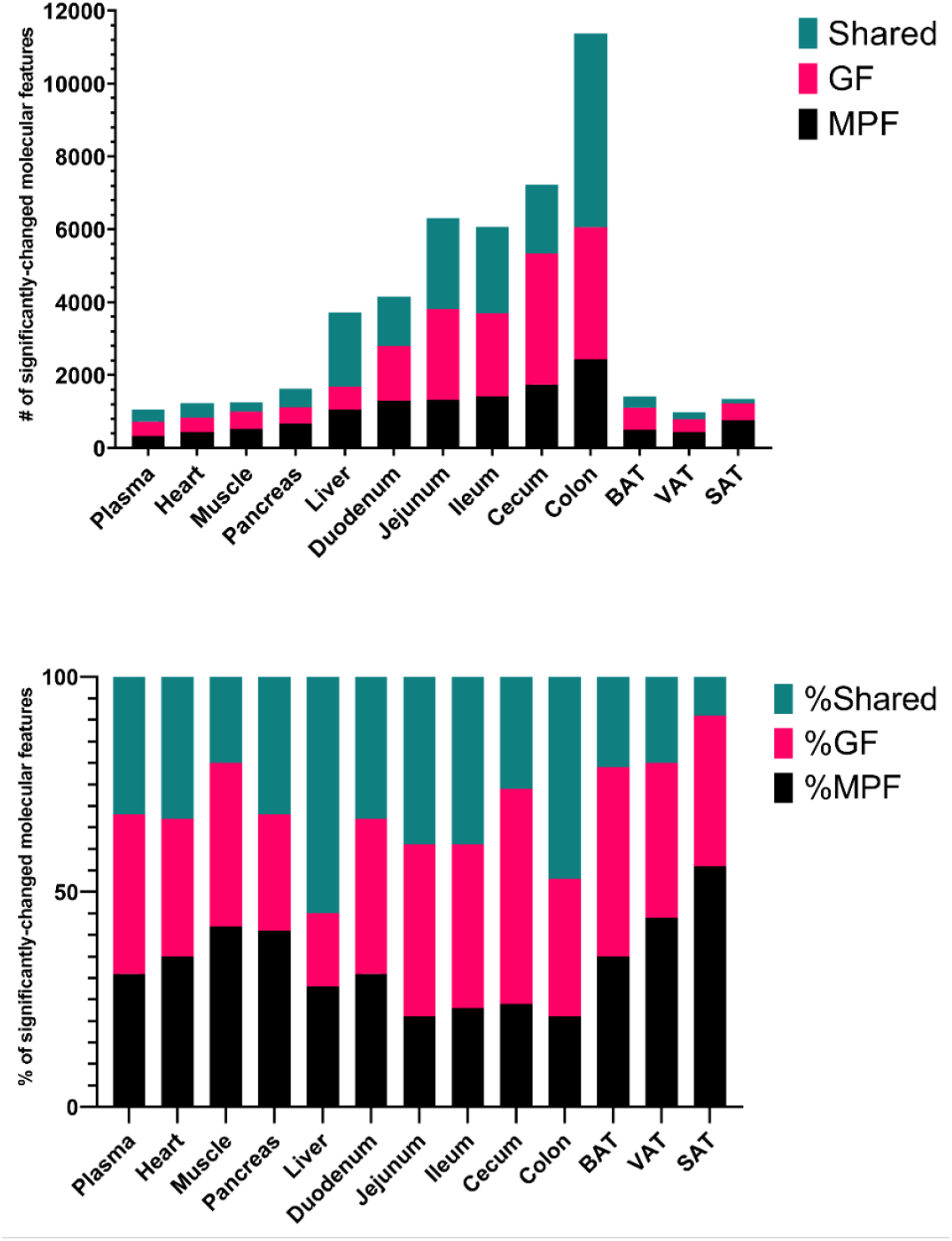
Number and percentage of significant molecular features per tissue. The total number of total significant molecular features from each tissue sourced from MPF only, GF only or shared (*Upper*), Percentage of significant unique molecular features from each murine class per tissue (*Lower*). Level of significance is defined as having a fold change ≥ 1.3, *p*-value ≤ 0.05, and *q*-value ≤ 0.05

To assess the differences in the metabolite composition in the GF and MPF mice, we performed PCA on the molecular features across all the tissues. The metabolic profile was influenced by the tissue type as the main driving factor as well as by the colonization status (*i.e.*, GF or MPF), indicating that the extensive effect of microbiota on the metabolite composition extends to peripheral organs (Figure 2).

**Figure 2.**
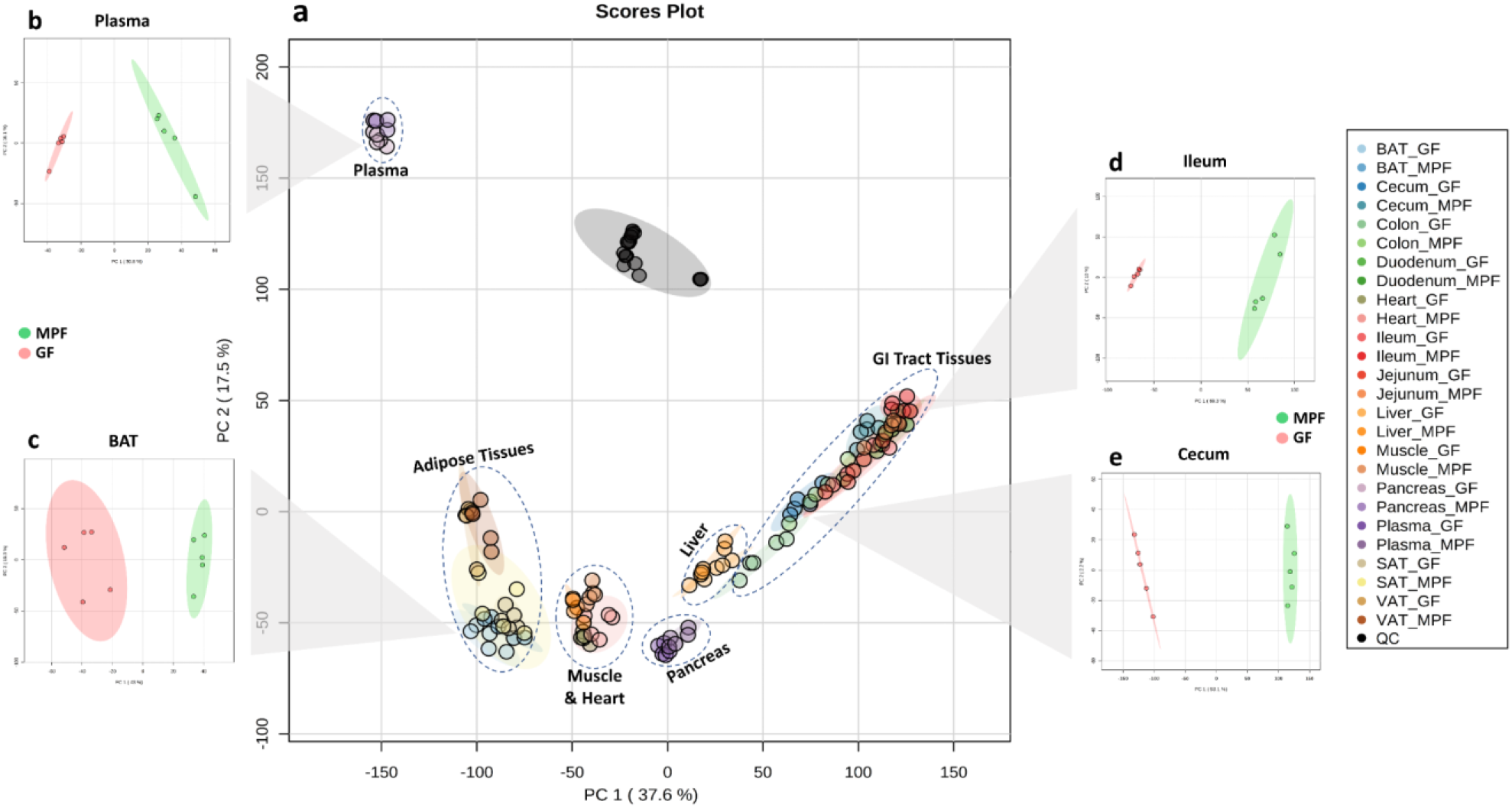
Principal component analysis (PCA) of the profiling data shows separation between tissues and mice groups. Data shown for reverse phase (RP) and HILIC modes with both positive and negative ionization **a)** PCA of all the analyzed samples from all tissues, **b)** PCA of plasma samples from the GF and MPF mice, **c)** PCA of BAT samples from the GF and MPF mice, **d)** PCA of ileal samples from the GF and MPF mice, **e)** PCA of cecal samples from the GF and MPF mice.

Further detailed investigation on the differential molecular features in each tissue, demonstrates how the magnitude of the impact of the colonization status varies across the different tissue types (Figure 3, Supplementary Figure 1). Likewise, on the level of individual metabolites and metabolite classes, it was evident that the variation across organs was high, as some of metabolites were differential only in one tissue and some others across several, for example tauro-alpha/beta-muricholic acid and *p*-cresol glucuronide were differential in multiple tissues and present only in GF and MPF mice respectively, whereas particular phospholipids were only differential in SAT and mostly present in GF mice (Figure 3 and Supplementary Figure 1).

**Figure 3.**
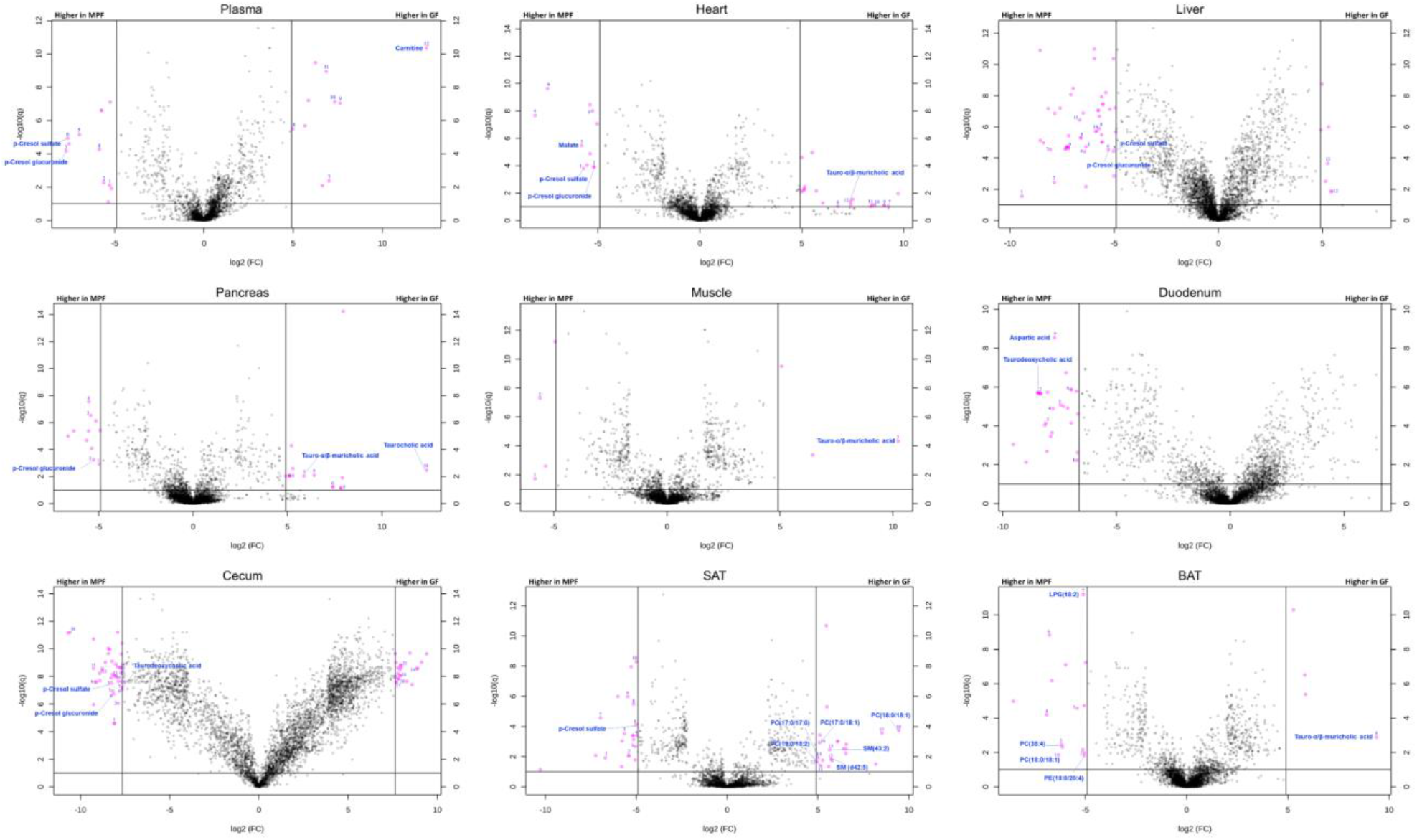
Volcano plots of the molecular features detected in nine representative tissues. The illustrated tissues include plasma, heart, liver, pancreas, muscle, duodenum, cecum, subcutaneous adipose tissue (SAT), and brown adipose tissue (BAT); see Supplementary Figure 1 for volcano plots of all studied tissues individually with selected metabolites annotated. The binary logarithm of the fold change (FC) is shown as the function of the negative common logarithm of the *q*-value (false discovery rate corrected *p*-value). A positive log_2_(FC) signifies a higher abundance in the GF mice compared to the MPF mice. The purple dots represent molecular features fulfilling the significance criteria (FC > 200 or FC < 0.005 for the cecum and the colon tissues, FC > 100 or FC < 0.01 for duodenum, jejunum, and ileum, FC > 30 or FC < 0.033 for the rest of the tissue types, and q < 0.1). Molecular features are presented as their binary logarithmic fold change [log2(FC)]against the negative common logarithm of the q-value [false discovery rate corrected p-value; −log10(q)] of the differential expression between the GF and MPF mouse group. Note: Although the purple dots represent molecular features fulfilling the above-mentioned significance criteria, and were unique to the sample type.

We further applied *k*-means cluster analysis to examine the abundance of the differential molecular features between the mice groups across all tissue types. In our data, the molecular features were clustered based on 1) their presence in the different tissues, 2) their presence in the GF or MPF mice (Supplementary Figure 2). Among the 10 clusters generated by *k*-means clustering analysis, clusters 2, 4, and 5 (1,214 molecular features in total) contained the metabolites that showed significant differences between the GF and MPF mice that were unique to one tissue or a subset of tissues. Cluster 2 indicates the subset of molecular features that were significantly higher in the cecum and the colon of the GF mice. Cluster 4 represents a subset of molecular features that were significantly higher in the cecum and the colon of the MPF mice. Cluster 5 shows the metabolites that were significantly higher in the duodenum, jejunum, ileum, and liver of the MPF mice. It is notable that no other clusters contained features that were either unique to the GF or MPF subset of a tissue. After implementing PCA, volcano plots, and *k*-mean cluster analyses, the differential molecular features across all tissue types were taken into examination for metabolite identification. Supplementary Table 2 lists the annotated metabolites organized into their respective chemical classes and metabolic pathways for the thirteen tissues and their sub-groups (GF and MPF). Figures 4–8 illustrate the annotated compounds across different metabolite classes in a heatmap chart, and will be discussed in following chapters.

**Figure 4.**
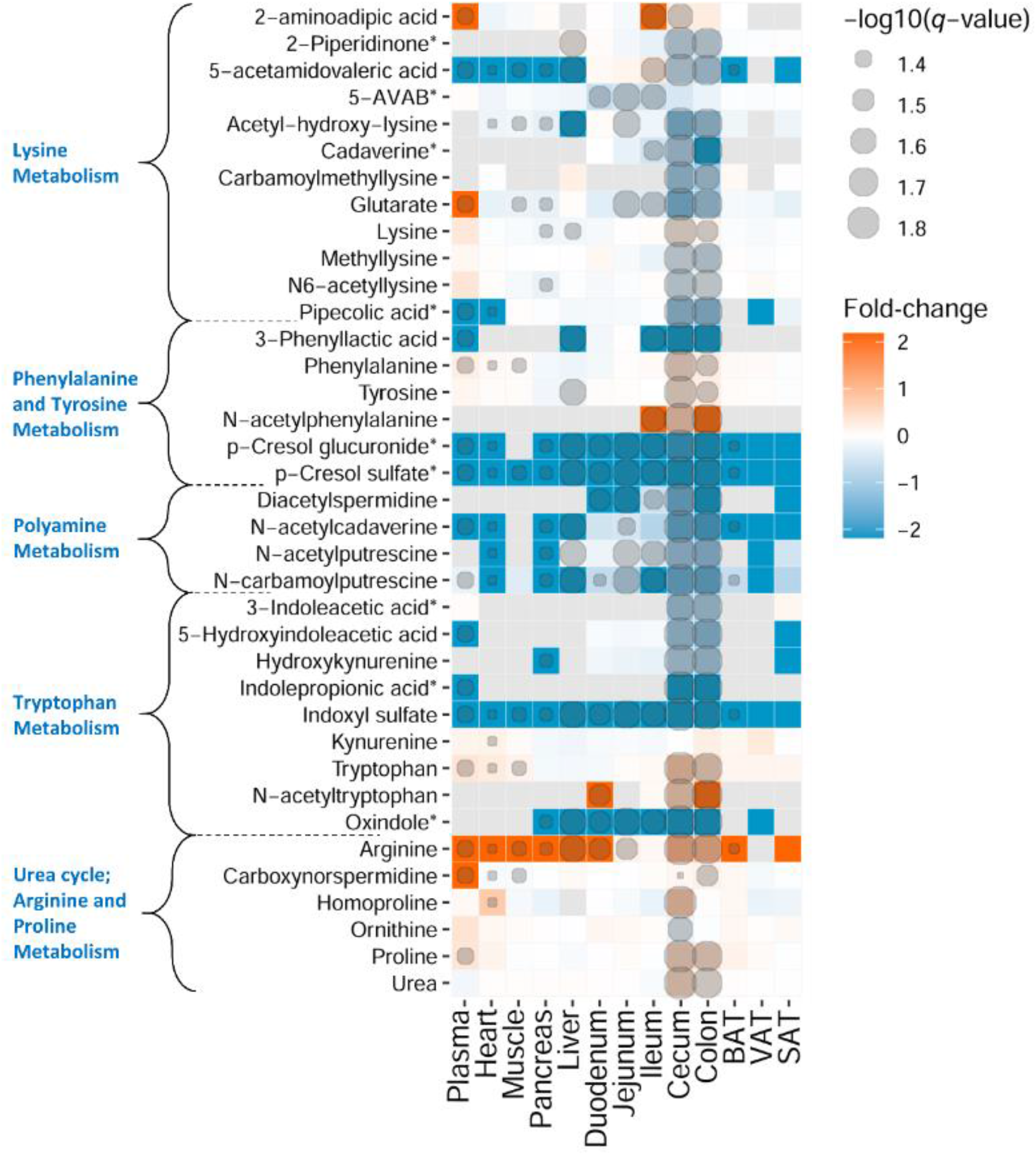
Heatmap representation of identified metabolites in amino acid chemical class. Fold-change and degree of significance comparisons were performed between the GF and MPF within each tissue (Mann–Whitney U-test and Benjamini and Hochberg false discovery rate correction *p*-value ≤0.05, and *q*-value ≤ 0.05). Each comparison for a tissue is represented by a colored cell. Gray cells represent metabolites that were not found in the tissue. <u>Orange and blue</u> cells represent metabolites more abundant in GF and MPF mice, respectively. *Metabolite is known to be bacterial-borne.

### 3.2. Microbiota affects various branches of amino acid and peptide metabolism

Our results show that majority of the amino acid metabolic pathways, including lysine, phenylalanine and tyrosine, polyamine, tryptophan, and urea cycle; arginine and proline pathways, were affected by the presence or absence of the microbiota (Figure 4). In lysine metabolic pathway, the level of lysine and 2-aminoadipic acid were higher in the GI tract tissues of GF mice than in their respective MPF group. In contrast, the rest of the identified metabolites, including N-acetyllysine, pipecolic acid, methyllysine, acetylhydroxylysine, carbamoylmethyllysine, glutarate, 5-acetamidovaleric acid, 5-aminovaleric acid betaine (5-AVAB), 2-piperidinone, cadaverine, and N-acetylcadaverine showed lower abundance in the GF mice. Additionally, 2-aminoadipic and glutarate were also higher in the plasma of GF mice.

In phenylalanine and tyrosine metabolic pathway, the levels of phenylalanine, tyrosine, and N-acetylphenylalanine were significantly higher in the GI tract of GF mice than in their respective MPF counterparts. Other identified phenylalanine and tyrosine metabolites, *p*-cresol glucuronide, *p*-cresol sulfate, and 3-phenyllactic acid had lower abundance in the GF mice (Figure 4). Notably, *p*-cresol glucuronide and *p*-cresol sulfate were found across all the examined tissue types, and as illustrated in the volcano plots (Figure 3), they were also the most differential metabolites in multiple tissues; *p*-cresol glucuronide in plasma, heart, liver, pancreas, cecum, and colon, *p*-cresol sulfate in plasma, heart, liver, cecum, and SAT.

In tryptophan metabolic pathway (indole-containing compounds), tryptophan was significantly higher in abundance in the plasma, heart, muscle, cecum, and colon of GF mice (Figure 4). The abundance of N-acetyltryptophan in the duodenum, cecum and colon of GF mice was also significantly higher than in their respective MPF counterparts. In contrast, abundance of tryptophan metabolites including 3-indoleacetic acid, 5-hydroxyindoleacetic acid, indolepropionic acid, hydroxykynurenine, and oxindole were significantly lower in the cecum and colon of GF animals. Additionally, 5-hydroxyindoleacetic acid, indolepropionic acid showed significantly low abundance in the plasma GF mice, and oxindole was significantly lower in all the GI tract tissues as well as in the pancreas and the liver.

All the identified metabolites from the polyamine metabolic pathway (N-acetylputrescine, N-carbamoylputrescine, and diacetylspermidine) were observed throughout various examined tissues excluding muscle, and had a lower abundance in the GF group in most of the tissues (Figure 4).

In the urea cycle; arginine and proline metabolic pathways, the identified metabolites, urea, arginine, proline, homoproline, and carboxynorspermidine were markedly more abundant in the GI tract of GF mice than in that of the MPF mice, whereas only ornithine showed lower abundance in the GI tract of GF mice. Interestingly, BAT was the only tissue having a higher abundance of all the identified metabolites from this metabolic pathway in the GF mice (Figure 4).

The results observed from the effect of the microbiota on peptide metabolism show a higher abundance of almost all the identified oligopeptides (di-, tri- and tetrapeptides) in the GI tract of GF mice (Figure 5). However, peptides were not among the most significantly changed metabolites across the different tissues, with the exception of glutamylglutamic acid (Glu-Glu) and aspartyltyrosine (Asp-Tyr) that were the two peptides were also shown to be significantly higher in the cecum tissue of GF from volcano plots (based on the inclusion criteria from volcano plots with molecular features above the threshold (Figure 3).

**Figure 5.**
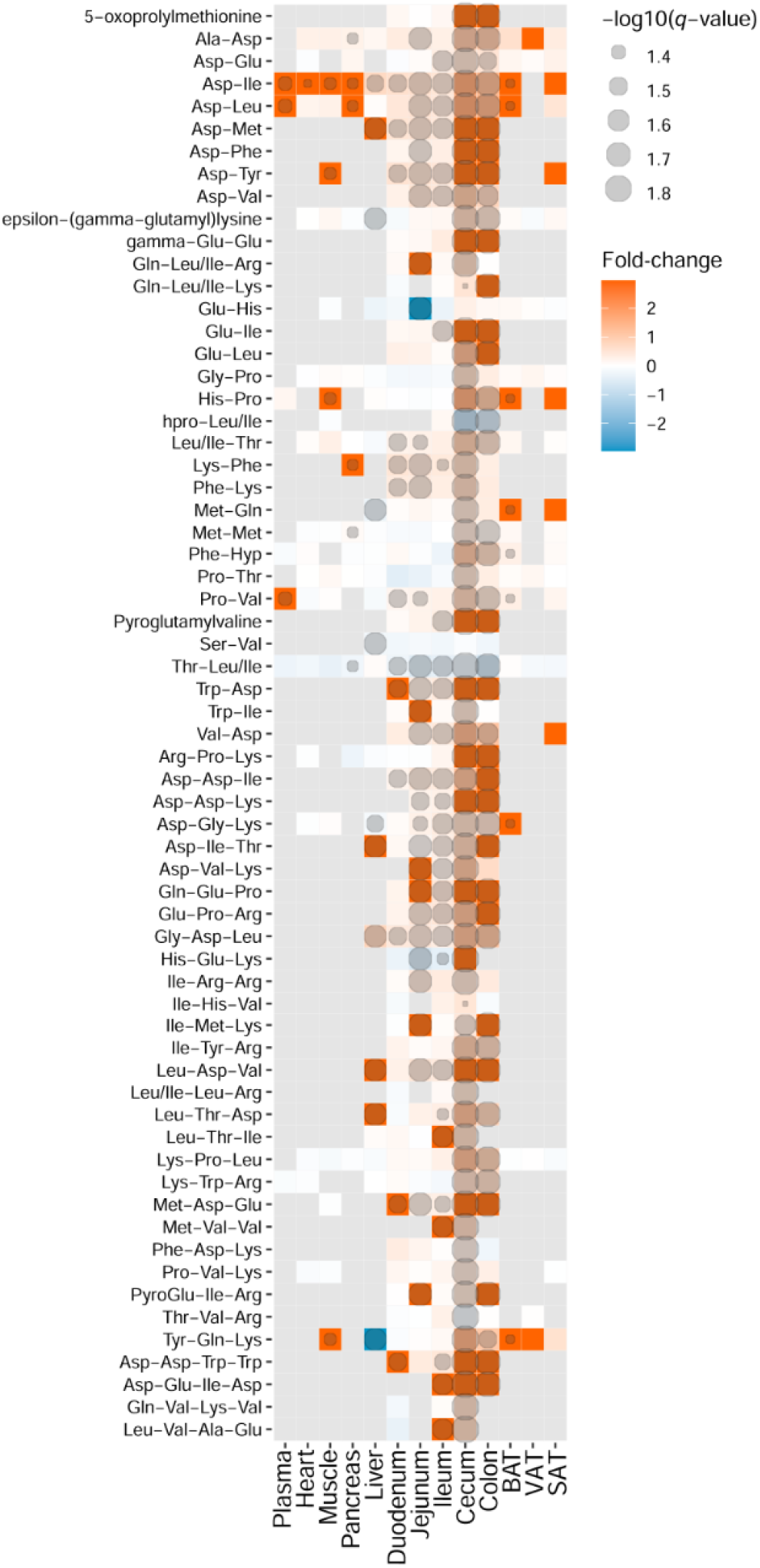
Heatmap representation of identified metabolites in peptide chemical class. Fold-change and degree of significance comparisons were performed between the GF and MPF within each tissue (Mann–Whitney U-test and Benjamini and Hochberg false discovery rate correction *p*-value ≤0.05, and *q*-value ≤ 0.05). Each comparison for a tissue is represented by a colored cell. Gray cells represent metabolites that were not found in the tissue. <u>Orange and blue</u> cells represent metabolites more abundant in GF and MPF mice, respectively.

### 3.3. Carbohydrates, products of energy and lipid metabolism are modified in response to the microbiota

The results herein indicated a higher level of mono- and disaccharides in the GF mice, particularly in the cecum and the colon (Figure 6). In this study, the abundance of the identified metabolites, aconitic acid, citric acid, isocitric acid, and malate from energy metabolism, was higher in the GI tract tissues of GF mice. All of the mentioned metabolites were lower in abundance in the heart tissue and malate was absent in the heart of GF mice (Figure 6). Notably, as also illustrated in the volcano plots (Figure 3), malate was among the most differential compounds in the heart of MPF mice.

**Figure 6.**
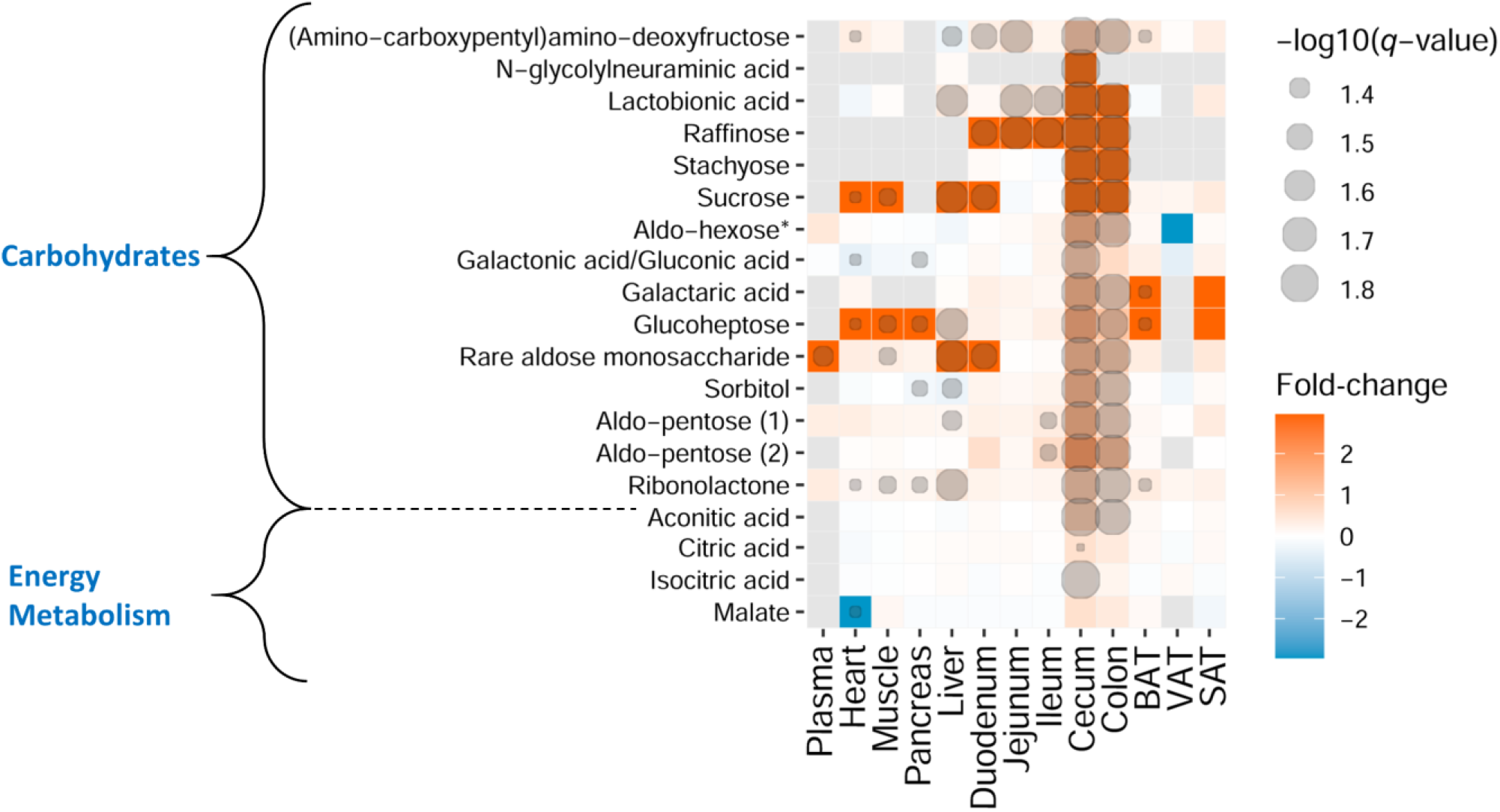
Heatmap representation of identified metabolites involved in carbohydrate and energy metabolism. Fold-change and degree of significance comparisons were performed between the GF and MPF within each tissue (Mann–Whitney U-test and Benjamini and Hochberg false discovery rate correction *p*-value ≤0.05, and *q*-value ≤ 0.05). Each comparison for a tissue is represented by a colored cell. Gray cells represent metabolites that were not found in the tissue. <u>Orange and blue</u> cells represent metabolites more abundant in GF and MPF mice, respectively. * This metabolite is one the following isomers: Mannitol, Galactitol, Iditol

Bile acids were among the most differential metabolites affected by the gut microbiota, and they were found not only in the GI tract and liver, but throughout the examined tissues including heart, pancreas, and fat tissues (Figure 7). Germ-free animals had lower levels of identified primary bile acids, including chenodeoxycholic acid, allocholic acid, β-muricholic acid, norlithocholic acid, three isomers of hydroxy-oxo-cholan-24-oic acid, 7-HOCA (7alpha-hydroxy-3-oxo-4-cholestenoate), cholanic acid-diol-sulfoethyl-amide, and 7-sulfocholic acid. In contrast, the identified conjugated primary bile acids, taurocholic acid, and one of its isomers (*i.e.*, taurallocholate, tauroursocholate, or taurohyocholate), tauro-α/β-muricholic acid, and several unidentified glycine and taurine-conjugated bile acids were higher in the GF mice (Figure 7). The increase of taurine conjugated bile acids in the GF status was particularly evident in e.g., VAT tissue (Supplementary Figure 1). The identified secondary bile acids, including 7-ketodeoxycholic acid, hyodeoxycholic acid, deoxycholic acid, taurochenodesoxycholic acid (taurochenodeoxycholic acid), taurodeoxycholic acid, lithocholic acid, and ursodeoxycholic acid were either completely missing from the GF mice or were present in negligible amounts.

**Figure 7.**
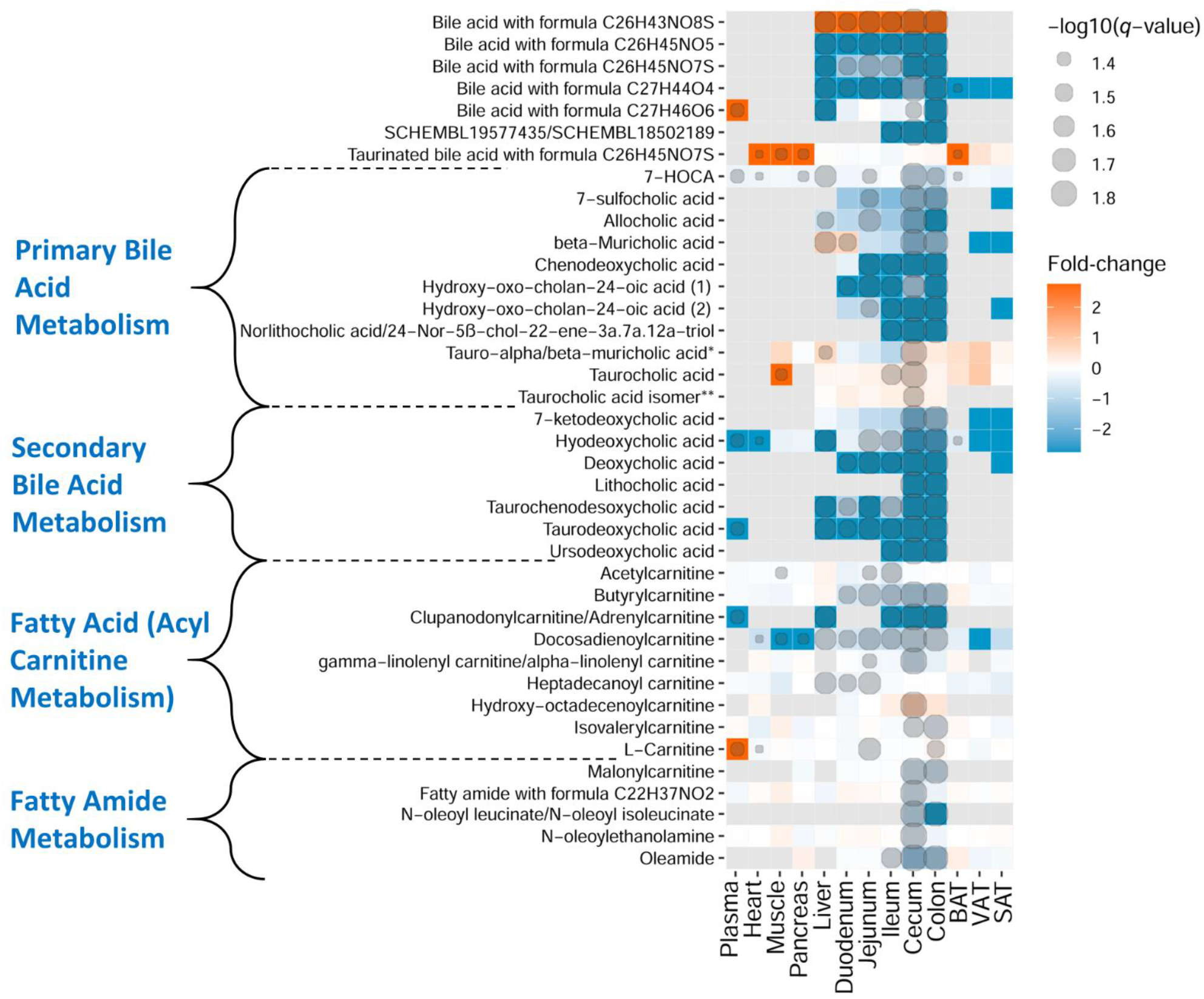
Heatmap representation of identified metabolites in bile acids, fatty amides, carnitine, and acylcarnitine metabolism classes. Fold-change and degree of significance comparisons were performed between the GF and MPF within each tissue (Mann–Whitney U-test and Benjamini and Hochberg false discovery rate correction *p*-value ≤0.05, and *q*-value ≤ 0.05). Each comparison for a tissue is represented by a colored cell. Gray cells represent metabolites that were not found in the tissue. <u>Orange and blue</u> cells represent metabolites more abundant in GF and MPF mice, respectively. *Despite comparing the spectra against the purified standard, we were not able to differentiate between tauro-α-muricholic acid and tauro-β-muricholic acid. **Taurocholic acid isomer: either taurallocholate or tauroursocholate or taurohyocholate.

In the carnitine and acylcarnitine metabolic pathways, carnitine, acetylcarnitine, clupanodonylcarnitine/adrenylcarnitine, malonylcarnitine, butyrylcarnitine, isovalerylcarnitine, linolenylcarnitine, and heptadecanoylcarnitine, showed lower abundance in the GI tract of GF mice, while hydroxyoctadecenoylcarnitine showed the higher abundance in the same mouse line compared to the MPF mice. Interestingly, L-carnitine had higher abundance in plasma samples of GF mice (Figure 7). All the detected fatty amides were low in abundance in the GF mice in the cecum and the colon (Figure 7).

### 3.4. Flavonoids, phenolic acid derivatives, and terpenes were other chemical classes influenced by microbiota

The intestinal microbiota plays an important role in the metabolism of plant-derived phytochemicals including flavonoids, phenolic acid derivatives, and terpenes. The results herein indicate that the differences in identified flavonoids, phenolic acid derivatives, and terpenes between the GF and MPF mice were mostly found in the GI tract, and these differences were higher in or exclusive to the GF animals with a few exceptions’ high abundances in the MPF mice (Figure 8). Exceptions included daidzein, 3-(3-hydroxyphenyl)propionic acid, dihydrocaffeic acid, gentisic acid, and medicagenic acid along with three unidentified flavonoids with molecular formulas C_18_H_18_O_8_ (two isomers) and C_21_H_22_O_10_ and one triterpenoid with formula C_30_H_48_O_2_ that were higher in GI tract of MPF mice. More specifically, gentisic acid, a bacterial end-metabolite of dietary (plant) salicylic acid [52], along with microbial degradation products of the dietary phenolic acids, including 3-(3-hydroxyphenyl)propionic acid and dihydrocaffeic acid, were absent or existed in low amounts in the GF mice tissues. Various derivatives of the isoflavonoid compounds including daidzein and its sulfated form, formononetin and its glucuronide, genistein glucuronide, and genistein sulfate were also annotated in the data, and the compounds were differential in GI tract of the animals. Notably, as also illustrated in the volcano plots (Figure 3), 3-(3-hydroxyphenyl)propionic acid was among the most differential metabolites in the cecum and the colon of MPF mice. Ferulic/isoferulic acid sulfate and vanillic acid sulfate were higher in the cecum of GF mice.

**Figure 8.**
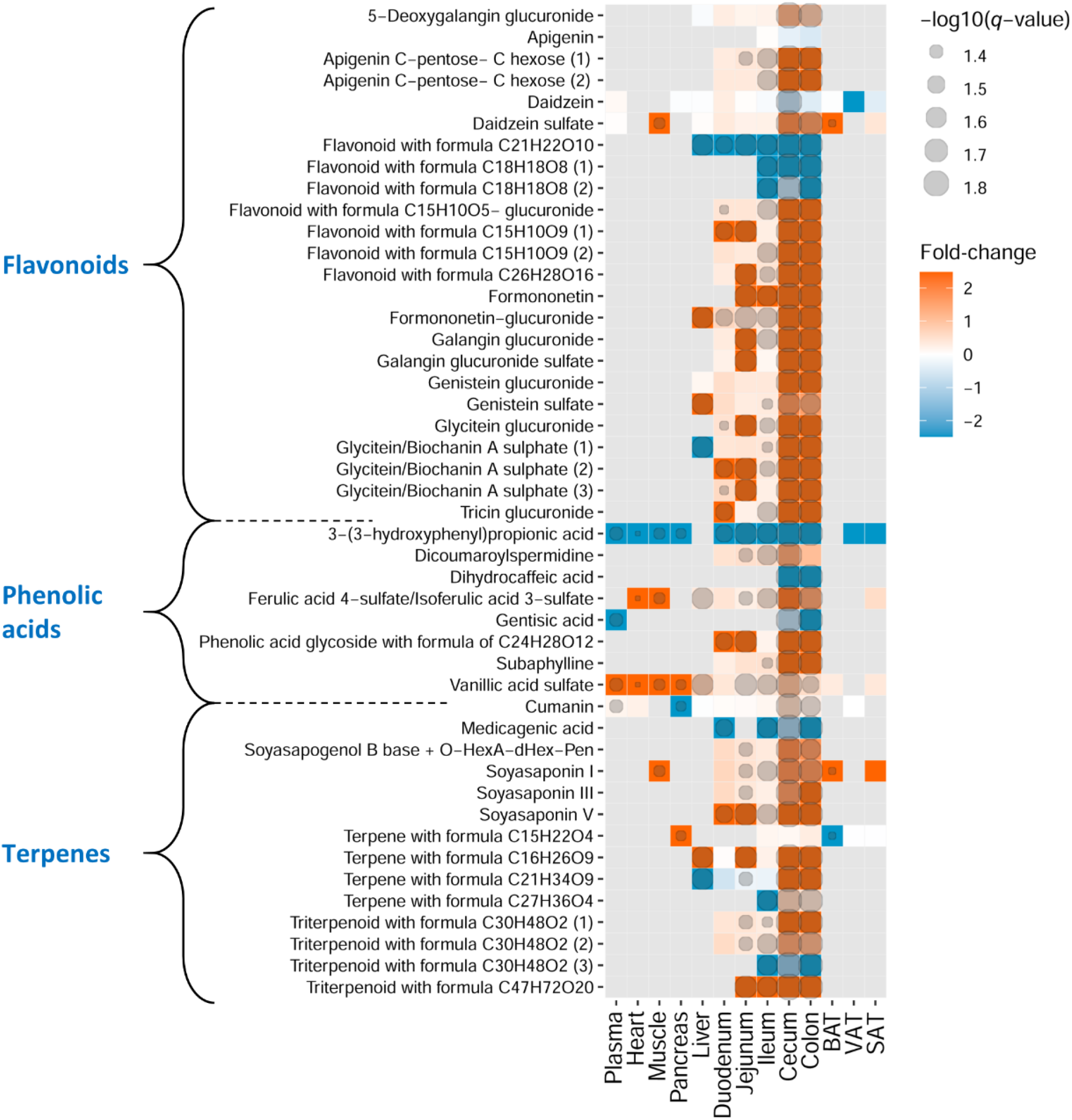
Heatmap representation of identified metabolites in phenolic acid derivatives, flavonoids, and terpenes. Fold-change and degree of significance comparisons were performed between the GF and MPF within each tissue (Mann–Whitney U-test and Benjamini and Hochberg false discovery rate correction *p*-value ≤0.05, and *q*-value ≤ 0.05). Each comparison for a tissue is represented by a colored cell. Gray cells represent metabolites that were not found in the tissue. <u>Orange and blue</u> cells represent metabolites more abundant in GF and MPF mice, respectively.

## 4. Discussion

In this study, we observed how the presence or absence of gut microbiota has a strong influence on the biochemistry of mammalian tissues and revealed substantial variation in the distribution of metabolites from distinct chemical classes. The most affected metabolite classes were amino acids, peptides, carbohydrates, metabolic products of energy metabolism, lipids particularly, bile acid, fatty amide, and acyl carnitine metabolism, and plant-derived phytochemicals, including flavonoids, phenolic acid derivatives, and terpenes. We herein also reported that all the GI tract tissues as well as plasma and BAT were the most affected tissues in the GF mice and showed higher percentage of significantly abundant metabolites when compared to their counterpart MPF mice in the same tissues.

Given the fact that the GF status of the mice has an impact on the number of the significant metabolites [53], this may suggest that with a lack of gut microbiota and, subsequently, its related compounds in the GF mice, the host endogenous metabolism gives rise to different metabolic pathways that compensate for the loss or absence of microbiota [54]. This hypothesis may also be supported by the high abundance of metabolites from energy metabolism in the GF mice. Our results established that multiple tissue metabolites are potentially derived from microbiota. Metabolic pathways, endogenous and diet-derived metabolites that were modulated by gut microbiota in the mice merit to be studied further for the potential synergistic health implications.

Proteins and amino acids are a main part of the diet. In addition to serving as nutrients, they play a crucial role in maintaining the gut microbiota and energy metabolism. For instance, some gut microbial species are able to yield energy from the oxidized forms of the branched-chain amino acids, which also leads to SCFAs as well as branched-chain fatty acids production [55]. It is also evident that gut bacteria play an important role in host amino acid homeostasis [56–58]; once taken up by bacteria, amino acids can be either incorporated into bacterial cells for protein biosynthesis, or become catabolized and used as energy source, or get biotransformed to a diverse range of bioactive molecules such as conversion of tryptophan to other indole-containing metabolites [59, 60]. Our results showed a significant presence of peptides, mostly in the GI tract of GF mice, most likely due to the absence of gut microbiota to hydrolyze them further. Likewise, lysine, urea, proline, arginine, and the aromatic amino acids tryptophan, phenylalanine, and tyrosine with their acetylated forms, N-acetyltryptophan and N-acetylphenylalanine serve as substrates for multiple pathways in gut microbiota, and all of these were indeed accumulating in the GI tract in the absence of gut microbiota [43, 59–61]. Thus, higher levels of these metabolites in the GI tract of GF mice may explain the critical role of gut microbiota in expanding the compound diversity. Indoleacetic acid, hydroxyindoleacetic acid, indolepropionic acid, and oxindole are microbial catabolites of tryptophan and existed in higher abundance in the cecum, colon, and some in the plasma as well in the MPF mice. Microbial tryptophan catabolites are shown to have potential role in mediating microbe-host interactions and eventually contribute to the health status of the host [62]. Indoleacetic acid, hydroxyindoleacetic acid, and indolepropionic acid are shown to regulate gut barrier function [62, 63] and merit further investigation for their potential role in reducing likelihood of cardiovascular diseases [64] and developing type 2 diabetes [65, 66] likely by modulating host metabolism through the production of glucagon-like peptide-1 (GLP-1) to improves insulin resistance [62]. Oxindole, another tryptophan microbial metabolite, was higher in all the GI tract tissues as well as in the liver and plasma in the MPF mice. High abundance of this metabolite is observed in impairment of insulin secretion in pancreatic beta cells and in hepatic encephalopathy conditions [67, 68] and merit further investigations for its potential role as a biomarker of hepatic cirrhosis. In phenylalanine and tyrosine metabolic pathways, *p*-cresol sulfate, *p*-cresol glucuronide, and 3-phenyllactic acid are known metabolites of gut microbiota and lower levels of these metabolites in the GF mice are expected [69–71]. *p*-cresol which is the precursor of *p*-cresol sulfate and *p*-cresol glucuronide, is one of the end product of tyrosine and phenylalanine biotransformation by intestinal bacteria [72]. *p*-cresol exert many biological and biochemical toxic effects and should be excreted from the body [72]. In one of the mechanisms when this methylphenol metabolite reaches the mucosa of the colon [73] and liver [74], sulfatation and glucuronidation take place, as a result *p*-cresol sulfate and *p*-cresol glucuronide are generated. The two latter metabolites are less toxic and more water-soluble compared to the parent compound, *p*-cresol, which makes it easier for the body to ger rid of them via urinary excretion.

Other affected metabolic pathways in the study were lysine and polyamine pathways. Although lysine was one of the main dietary constituents of both mice groups in the study, the higher abundance of 2-aminoadipic acid in plasma, ileum, cecum, and colon of the GF mice may be due to lysine degradation by the host, as it is the key intermediate metabolite of lysine catabolism. Lower levels of N-acetyllysine, pipecolic acid, methyllysine, acetylhydroxylysine, 5-acetamidovaleric acid, 5-AVAB, 2-piperidinone, cadaverine, N-acetylcadaverine, and glutarate from lysine pathway in the GF mice may be explained by the lack of gut microbiota, as the bacterial catabolism of lysine is shown to be one of the main contributing factors responsible for the production of these metabolites [75, 76]. A majority of the foodstuff contains polyamines and microbiota produces these compounds are absorbed in great amounts in the large intestine [77]. The bacterial-derived sources of polyamines may be the reasons why the high abundance of these compounds was observed in the MPF mice in our study [78].

Even though conventional mice have been shown to synthesize more urea compared to germ-free ones [79], we observed elevated level of urea in the GF mice, particularly, in the cecum and colon. That is most likely because of the breakdown of urea by gut microbiota urease and its bioconversion into ammonia and carbon dioxide (Figure 9) [79]. This phenomenon may be the reason for higher levels of arginine and proline in the GF mice. Lower levels of ornithine in the same mouse group could be explained by the promotion of bacterial ornithine production from arginine, which was strongly accumulating throughout majority of tissues examined from the GF mice [80] as well as the bacterial inhibition of arginine biosynthesis from ornithine [81]. Higher abundance of carboxynorspermidine, another metabolite of the urea cycle; arginine and proline metabolism, may be explained by lack of microbiota in this mouse group to use this metabolite as a substrate for the biosynthesis of norspermidine [82].

**Figure 9.**
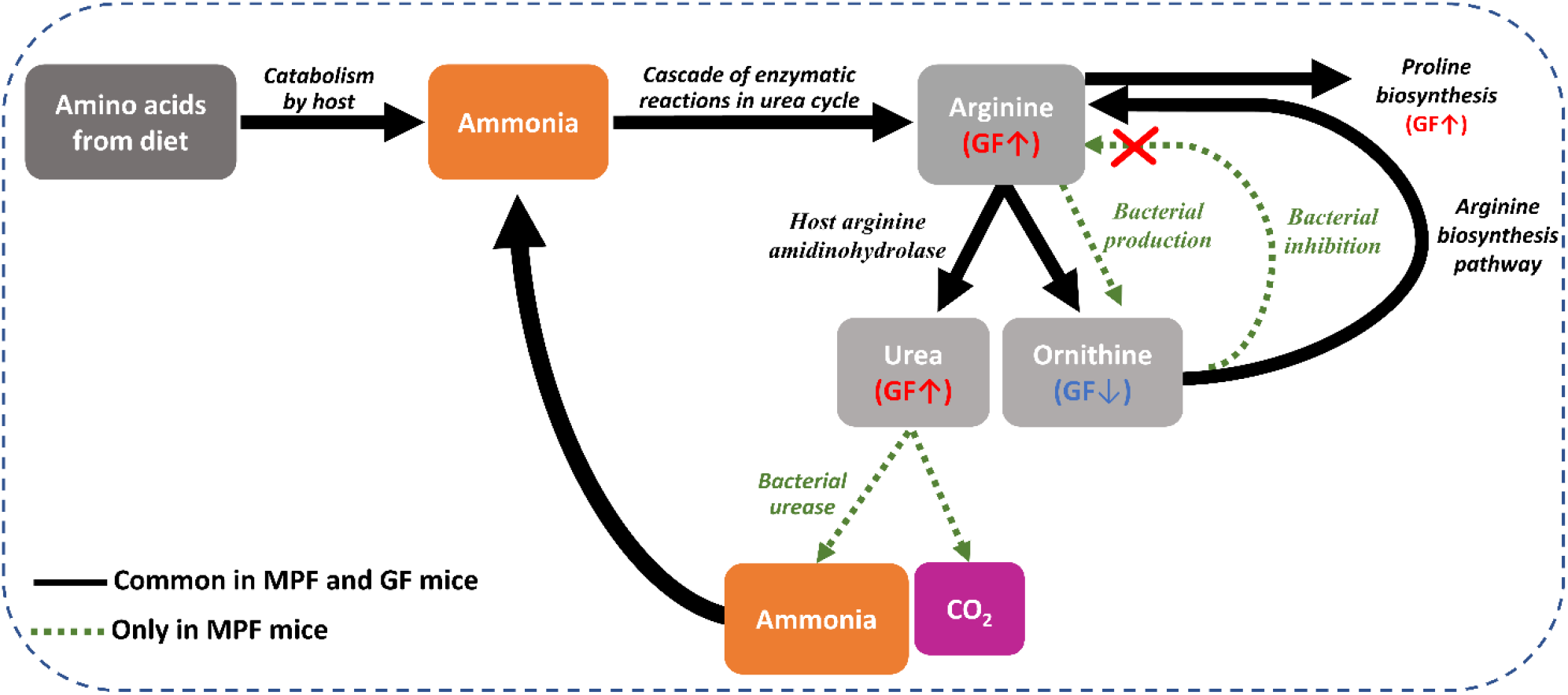
Summary of the fate of key metabolites (arginine, proline, urea, and ornithine) in the urea cycle and arginine and proline metabolism. Catabolism of dietary amino acids leads to the production of ammonia. Ammonia further undergoes conversion to urea via the urea cycle. In mammals with conventional gut microbiota, urea can be broken down to ammonia and CO_2_ by bacterial urease. The ammonia produced by microbiota is released into the GI tract and is taken up by host cells and serves as a substrate to synthesize arginine in the urea cycle. Within the urea cycle, arginine is then converted into urea and ornithine. Simultaneously, ornithine can also be synthesized by gut bacteria. Ornithine produced from the two mentioned pathways can enter the arginine biosynthesis pathway to synthesize more arginine; nevertheless, the bacterial inhibition of arginine biosynthesis can be inhibited by some bacteria. This excess amount of arginine can enter either the arginine and proline metabolism or the urea cycle.

Gut microbiota is shown to modulate the energy balance of the host by allowing the host to harvest energy more efficiently from the digested food [83]. Carbohydrates undergo fermentation by gut bacteria that leads to the production of SCFAs, which serve as a major source of energy for the gut bacteria as well as host intestinal epithelial cells [84]. Since the GF mice lack microbiota, the intestinal epithelial cells utilize sugars as the primary source of energy. As a result, a higher abundance of carbohydrates (mono- and disaccharides) in all the tissues from the GI tract of GF animals was observed when compared to the conventional ones. The contribution of gut microbiota to the energy homeostasis of the host is well-established [85]. Germ-free mice have been reported to be leaner than conventionally raised mice and they are protected against diet-induced obesity [86]. High abundance of aconitic acid, citric acid, isocitric acid, and malate from energy metabolic pathway in the GF mice can be explained by the lack of microbe-mediated increase in energy uptake. Therefore, the host intrinsic metabolism must compensate to maintain its energy homeostasis through upregulating its energy metabolism [83].

Among the lipid chemical class, the metabolic pathways that were significantly altered included bile acid metabolism, fatty amide metabolism, and fatty acid (acylcarnitine) metabolism. Gut microbiota and its composition can substantially influence the dietary fats and, consequently, the lipid metabolism in the host. It is shown that GF mice tend to be resistant to the metabolic changes after a high-fat diet, suggesting that gut microbes are important mediators of lipid-induced metabolic dysfunction [87]. Amongst different lipid subclasses, bile acids have gained special attention in their relation to gut microbiota because of secondary bile acids, which have a microbial origin [88]. Bile acids are crucial in aiding lipid digestion and as dynamic signaling molecules to regulate energy homeostasis, glucose metabolism, and innate immunity [89]. Likewise, in our study the bile acids were among the most widely distributed metabolites across the different tissue types, reflecting potentially-significant role in metabolism of different organs [90].The results herein showed lower levels of primary bile acids in the GI tract of GF mice. The bile acid metabolism occurring in the germ-free mice are rather complex and previous studies indicated high variability in bile acid metabolism between germ-free and conventional mouse models; the phenotypes conceivably depend on diet, genetic background, and the respective composition of the microbiota in the conventional control groups [91]. The profound modulatory impact of gut microbiota on bile acids (*i.e.*, deconjugation, dehydrogenation, and dihydroxylation of primary bile acids) is well-established. Mice lacking intestinal bacteria have no deconjugation effect on the amino acid moiety of conjugated primary bile acids, and as they pass through the GI tract, they remain intact, which was also suggested in our study as the higher abundance of tauro-conjugated primary bile acids were found in the GI tract of the GF mice. On the other hand, all the identified secondary bile acids were missing from the GF mice, confirming that gut microbiota was absent in these mice.

Another lipid subclass with a significant difference between the two mouse lines was fatty amides. Fatty amide metabolism may be interrelated with bile acid metabolism; in our data, the MPF mice showed a lower abundance of fatty amides in the upper part of the GI tract compared to the lower part. Conversely, we observed a higher abundance of tauro-conjugated bile acids in the upper part of the GI tract when compared to the lower part of the GI tract of the same mouse group. Some bile acids, including tauro-α- and β-muricholic acids (in mice) and ursodeoxycholic acid (in humans) are known to be potent antagonists of the bile-acid-activated nuclear receptor, farnesoid X receptor (FXR) [20, 92]. Additionally, in a study reported by Gonzalez *et al*., it was shown that FXR could be bound by a number of endogenous bile acids, including tauro-α- and β-muricholic acids in the upper section of the GI tract expected [93]. Given this fact, we herein speculate that because of this affinity between FXR and tauro-conjugated-muricholic acids, the fatty amides gene expression towards their biosynthesis is downregulated in the upper part of the GI tract of MPF mice. The low abundance of tauro-α- and β-muricholic acids in the lower part of the GI tract of the same mouse group may be justified by the gut microbiota enzymatic activity through deconjugation as they pass through the GI tract [93]. Therefore, fewer tauro-α- and β-muricholic acids are available to bind to FXR in the lower section of the GI tract. As a result, the FXR expression and fatty amide biosynthesis are upregulated in this area, and that may explain the higher abundances of fatty amides in the lower part of the GI tract of MPF mice (Figure 10). However, more experimentations such as gene expression analysis in conventional and germ-free mice are required to fully validate this proposed model.

**Figure 10.**
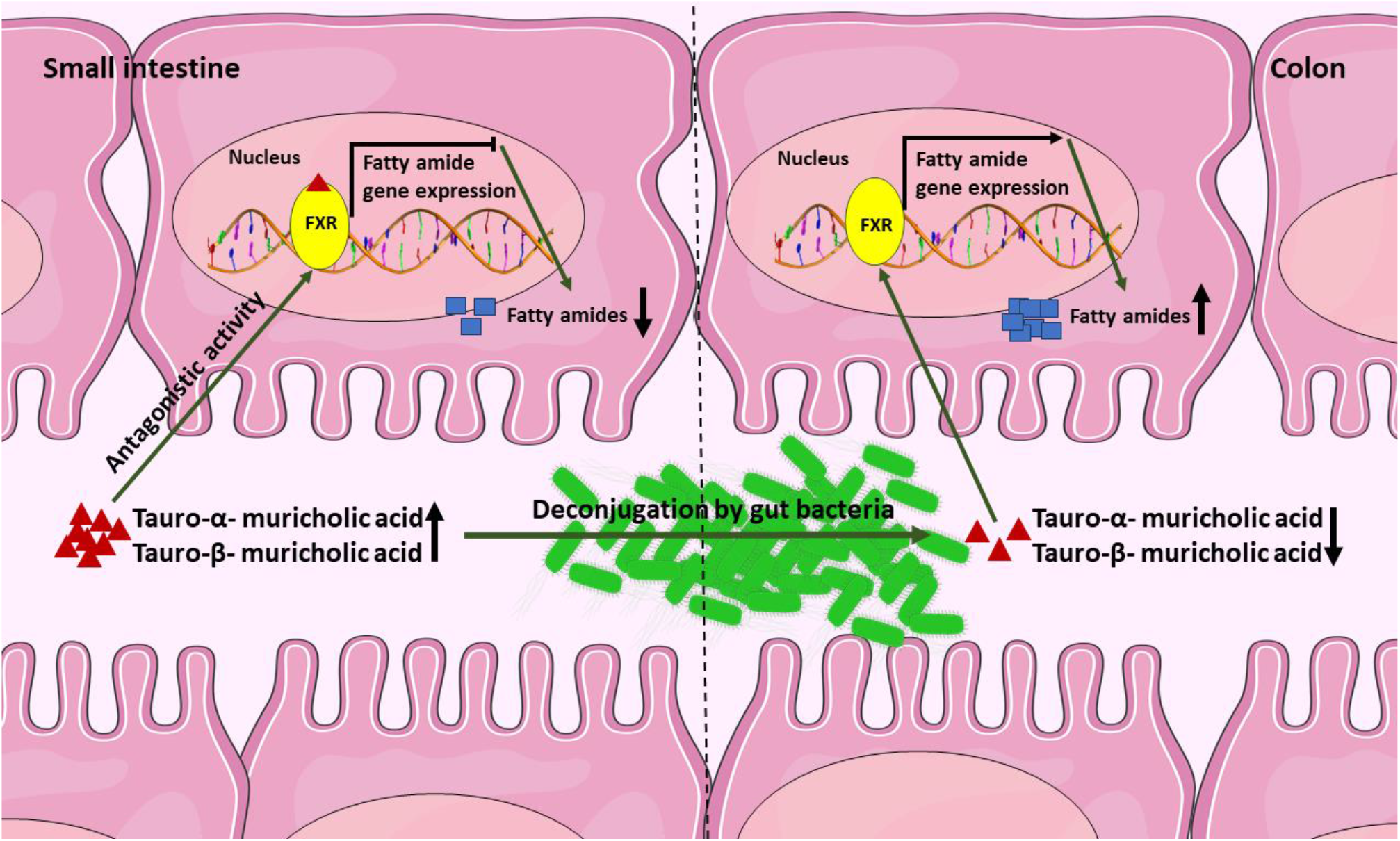
Proposed interrelation between tauro-conjugated muricholic acids and fatty amide biosynthesis in proximal and distal GI tract. When the FXR is bound to tauro-α- and β-muricholic acids in the upper section of the GI tract, the fatty amides biosynthesis is downregulated. As tauro-α- and -β-muricholic acids pass through the GI tract, they get deconjugated by gut microbiota. Therefore, there are fewer tauro-α- and β-muricholic acids are available to bind to FXR in the lower section of the GI tract. Thus, the FXR expression is upregulated in this area, and that may explain the higher abundances of fatty amides in the lower part of the GI tract.

Acylcarnitines are essential for oxidative catabolism of fatty acids as well as energy homeostasis in the host. Their accumulation in different organs is an indication of incomplete mitochondrial fatty acid β-oxidation and are important diagnostic markers of mitochondrial fatty acid β-oxidation disorders [94]. Some fatty acids can be converted to intermediate metabolites by gut bacteria. These bacterial fatty acids intermediate metabolites can further be absorbed and metabolized through β-oxidation in the host mitochondria, which then be converted into acylcarnitines and excreted (Supplementary Figure 3) [95]. This phenomenon may account for the role of gut microbiota in inducing acylcarnitine production in the MPF mice, as it was observed in this study, and that may be an indication of diminished fatty acid oxidation in germ-free mice. Amongst the metabolites that were present in the plasma, L-carnitine was the only detected metabolite from acylcarnitine metabolism pathway completely missing in plasma of MPF mice; aside from its main source, diet, L-carnitine is synthesized endogenously in liver and kidney from two essential amino acids, lysine and methionine. On the other hand, L-carnitine is metabolized by gut bacteria and liver in MPF mice to produce TMAO [96]. TMAO was our reference metabolite to assure the germ-free status of the mice used in the study, as it is a metabolite with microbial origin [46]. Besides the slow turnover of carnitine in the body [97], the absence of microbiota and lysine supplementation in the diet may account for higher plasma level of L-carnitine in GF mice. As mentioned, with the absence of microbiota, the GF mice are not able to convert carnitine to TMAO, and with additional lysine supplementation, this amino acid can enter the carnitine biosynthesis pathway to produce more carnitine.

In addition to dietary fibers, polyphenols, flavonoids, and terpenes are good examples of food components that favor the growth of probiotic bacteria in the colon [98]. Most of these compounds have very low bioavailability, and they reach the lower part of the GI tract wherein resident gut microbiota catabolizes them into smaller phenolic metabolites that are better absorbed and can have different biological effects using their unique hydrolytic enzymes (e.g., rhamnosidases). NIH #31M Rodent Diet comprises soybean and whole grains (wheat, oats, and corn). Therefore, it is likely that it contains a high number of phytochemicals that consequently reflect the mice tissue metabolome. In this study, although many of the identified phenolic acid derivatives flavonoids, isoflavonoids, and terpenes come from the diet, they remained intact in the GI tract of GF mice while they were absent or existed in low abundance in the GI tract of MPF mice. Interestingly, the identified metabolites from these chemical classes were mostly present in the GI tract of the animals. This may suggest that after microbial biotransformation, the transformed metabolites are more easily absorbed in the intestine and could exhibit enhanced bioavailability compared to their parent compounds [99]. Examples from this study include a higher abundance of daidzein and 3-(3-hydroxyphenyl)propionic acid in the majority of the tissues and dihydrocaffeic acid, gentisic acid, and medicagenic acid along with three unidentified flavonoids with molecular formulas C_18_H_18_O_8_ (two isomers) and C_21_H_22_O_10_ and one Triterpenoid with formula C_30_H_48_O_2_ in the cecum and the colon in MPF mice. Daidzein, the aglycone form of daidzin, is one of the widely studied isoflavones from soybeans. Following ingestion, daidzin is hydrolyzed by gut microbial glucosidases (some strains of *Bifidobacterium*, *Lactobacillus*, *Lactococcus,* and *Enterococcus),* which release the more bioavailable and potent metabolite against oxidative stress and cancer, daidzein [100, 101]. Acute and chronic exposure to daidzein in rats, has been associated with reduction of food intake and low adiponectin levels. Among various physiological properties that daidzein may have, it is known to be involved in lipid biosynthesis and that may suggest one of the reasons behind why daidzein levels exists in higher levels in adipose tissues of MPF mice tissue [102]. 3-(3-hydroxyphenyl)propionic acid and dihydrocaffeic acid are respectively the dehydroxylated and hydrogenated forms of caffeic acid (a common dietary component found in a variety of plant-derived food products) by gut microbiota (*Bifidobacterium*, *Escherichia*, *Lactobacillus*, and *Clostridium*) and merit attention for their antioxidant activities [103–106]. Gentisic acid, an active microbial metabolite of salicylic acid hydroxylation, is shown to inhibit colorectal cancer cell growth [52]. Some dietary phytochemicals such as ferulic/isoferulic acid and vanillic acid may be metabolized by gut microbiota. On the other hand, a higher abundance of their liver-modified metabolites ferulic/isoferulic acid sulfate and vanillic acid sulfate in the majority of the GF mice tissues may suggest that these latter metabolites escaped the microbial biotransformation, as they may be already absorbed in the upper intestinal track, and reached the liver to be converted into the sulfated form [107]. In general, while these characteristics of most of the identified phytochemicals and their derivatives that existed in higher abundance in the GF mice cannot be assessed in this study, it can be hypothesized that the presence of microbiota is essential in modulating the pool of phytochemicals entering the host tissues and exerting their physiological effect.

## 5. Study limitations

The important study limitations to consider in these outcomes was that with the current MS/MS databases, only a relatively small portion of the metabolites can be identified or putatively annotated. Additionally, in this study, the number of molecular features detected especially in the tissue samples, was remarkably high, and therefore the identification could only be focused on the fraction of the most differential features within each tissue. It is evident that the germ-free status caused massive differences across all the examined tissues and should therefore be addressed in a tissue-specific manner, alongside with other analytical techniques e.g., gene expression analysis. Models including germ-free animal studies have been widely used as a source of knowledge on the gut microbiota contributions to host homeostatic controls as they allow disruption in the gut microbiota to be studied in a controlled experimental setup [46]. However, when translating the results from gut microbiome research from mouse models to humans, there are pitfalls to be considered as these two species are different from another in anatomy, genetics and physiology [108]. Additionally, germ-free animals may not represent the best translational model for studying the functionality of the microbiota since they have intrinsically underdeveloped immunological responses, shortened microvilli, and many other structural and functional differences compared to conventional animals. However, studies showed that germ-free animals are valid *in vivo* experimental models for preclinical studies and for investigating the host-microbial interactions in health and diseases. Nevertheless, further investigations are needed to understand the massive impact microbiota on biochemicals at the interphase of food and human metabolism, and eventually, on health.

## 6. Conclusions

In summary, we have demonstrated a significant effect of the microbiome on the metabolic profile of 13 different murine tissues by applying a non-targeted metabolomics approach to a germ-free mouse model system. The results support the hypothesis that the chemistry of all major tissues and tissue systems are affected by the presence or absence of a microbiome. The strongest signatures come from the gut through the modification of host amino acids and peptides, carbohydrates, energy metabolism, bile acids, acylcarnitines, fatty amides, and xenobiotics, particularly the breakdown of plant-based natural products, flavonoids, and terpenes from food. Growing evidence from animal models and human studies supports that the microbiota is a key to various aspects of our health. As the link between humans and their microbial symbionts becomes more and more apparent, a combination of global non-targeted approaches and the development of tools that connect these data sets warrants greater attention to enable us to identify novel metabolites, leading to a better understanding of the deep metabolic connection between our microbiota and our health – with diet in between. We propose further investigations to uncover unique organ-specific microbial signatures and whether the specific inter-organ microbial signature can be linked to the host metabolic diseases such as obesity and its related disorders.

## Acknowledgments

The authors would like to thank Biocenter Finland (www.biocenter.fi) and Biocenter Kuopio (www.uef.fi/web/bck) for supporting the study.

## Abbreviations

GF: germ-free
MPF: murine-pathogen-free/conventional
GI: gastrointestinal
MAC: microbiota-accessible carbohydrate
SCFA: short-chain fatty acid
IBD: inflammatory bowel disease
VAT: visceral adipose tissue
SAT: subcutaneous adipose tissue
BAT: brown adipose tissue
TMAO: trimethylamine N-oxide
CAN: acetonitrile
HILIC: hydrophilic interaction chromatography
RP: reversed-phase
ESI: electrospray ionization
QC: quality control
LC-MS: liquid chromatography-mass spectrometry
MoNA: MassBank of North America
HMDB: human metabolome database
FC: fold change
FDR: False discovery rate
MeV: Multi experiment Viewer
PCA: principal component analysis
5-AVAB: 5-aminovaleric acid betaine
Glu-Glu: glutamylglutamic acid
Asp-Tyr: aspartyltyrosine
7-HOCA: 7alpha-hydroxy-3-oxo-4-cholestenoate
GLP-1: glucagon-like peptide-1
FXR: farnesoid X receptor

---

**Supplementary Figure 1.** Volcano plots of the molecular features detected in all the 13 representative tissues (ppt format).

**Supplementary Figure 2.**
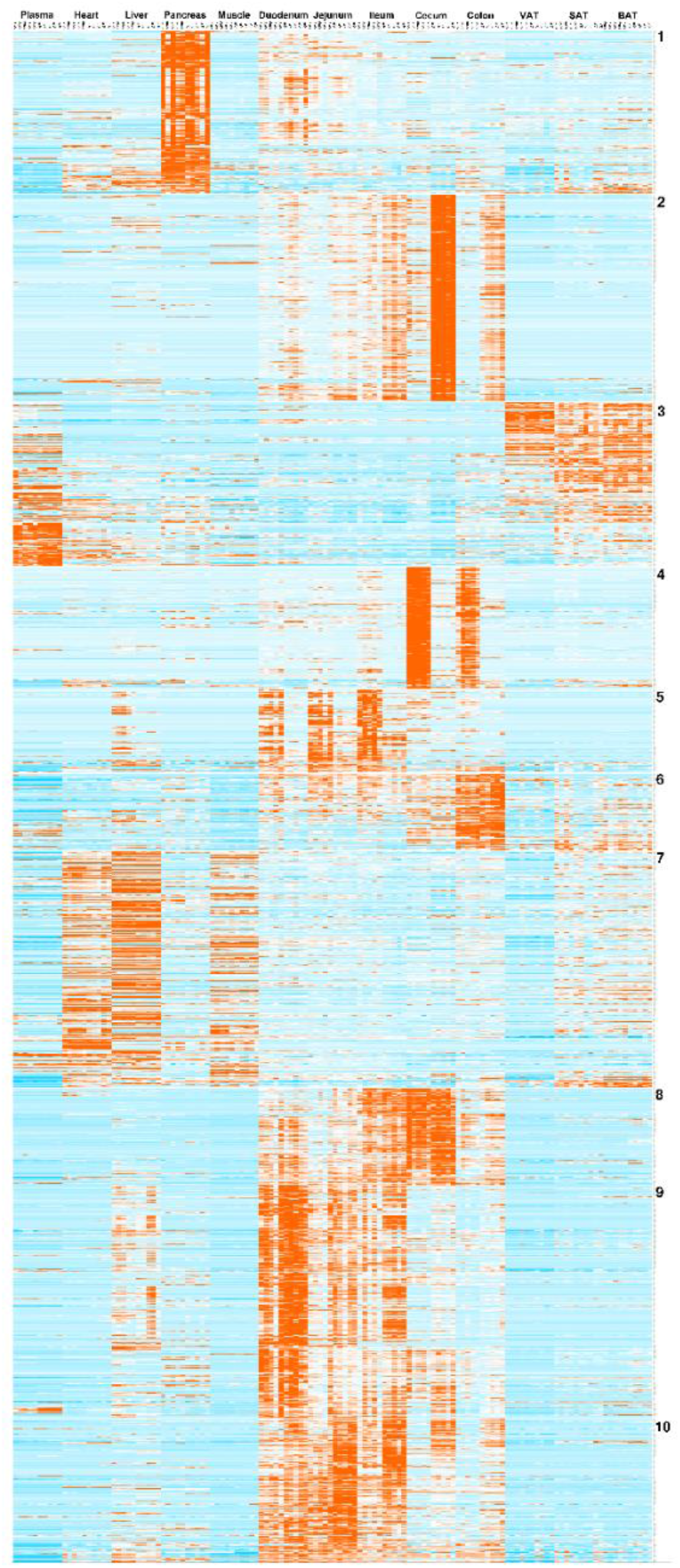
*k*-means cluster analysis of the filtered* molecular features (4,605 molecular features). Clusters 2, 4, and 5 containing 1,214 molecular features were considered for compound identification. Clusters 2 and 4 showed the molecular features that were mostly differential in the cecum and the colon and were higher in abundance in the GF and MPF mice, respectively. Cluster 5 showed the molecular features that were mostly differential in the duodenum, jejunum, ileum, and liver and were higher in abundance in the MPF mice. *Filtering criteria for inclusion were (1) p-value ≤0.05, and q-value ≤ 0.05, (2) high-intensity metabolite values (raw abundance ≥ 100,000), (3) containment of MS/MS fragmentation, (4) and retention time ≥0.7 min. For a molecular feature to be included, these inclusion criteria should exist at least in one tissue and one mouse group (GF or MPF). The different mouse groups are illustrated as follows with 5 mice in each group: Plasma MPF, Plasma GF, Heart MPF, Heart GF, Liver MPF, Liver GF, Pancreas MPF, Pancreas GF, Muscle MPF, Muscle GF, Duodenum MPF, Duodenum GF, Jejunum MPF, Jejunum GF, Ileum MPF, Ileum GF, Cecum MPF, Cecum GF, Colon MPF, Colon GF, VAT MPF, VAT GF, SAT MPF, SAT GF, BAT MPF, and BAT GF.

**Supplementary Figure 3.**
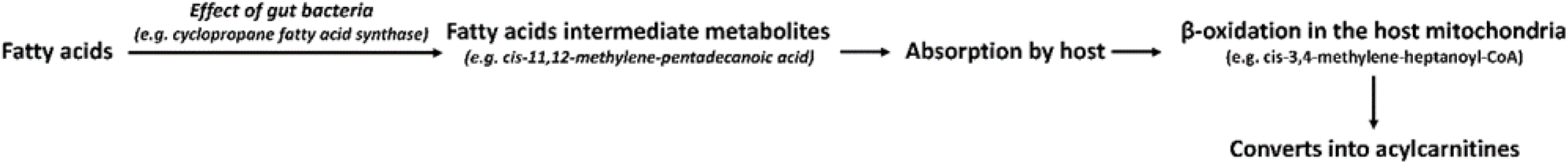
Induction of acylcarnitine production in the MPF mice (a suggested mechanism).

**Supplementary Table 1.** Summary of detected molecular features in all tissues from all the ionization modes.

| **Tissue** | **Total detected molecular features<sup>#</sup>** | **Total significantly changed molecular features\*** | **Significant change MPF > GF<sup>†</sup>** | **Significant change GF > MPF<sup>†</sup>** | **Unique to MPF<sup>†</sup>** | **Unique to GF<sup>†</sup>** | **Significant Shared<sup>†</sup>** |
| --- | --- | --- | --- | --- | --- | --- | --- |
| **Cecum** | 17894 (74%) | 11375 (64%) | 4603 (40%) | 6772 (60%) | 2438 (21%) | 3621 (32%) | 5315 (47%) |
| **Ileum** | 16659 (69%) | 6074 (36%) | 2198 (36%) | 3876 (64%) | 1419 (23%) | 2271 (38%) | 2380 (39%) |
| **Duodenum** | 16046 (66%) | 4149 (26%) | 1713 (41%) | 2436 (59%) | 1299 (31%) | 1505 (36%) | 1342 (33%) |
| **Jejunum** | 15613 (64%) | 6247 (40%) | 2257 (36%) | 3990 (64%) | 1321 (21%) | 2489 (40%) | 2500 (40%) |
| **Colon** | 15004 (62%) | 7226 (48%) | 2448 (34%) | 4778 (66%) | 1737 (24%) | 3606 (50%) | 1880 (26%) |
| **Liver** | 10021 (41%) | 3715 (37%) | 2295 (62%) | 1399 (38%) | 1055 (28%) | 628 (17%) | 2032 (55%) |
| **Pancreas** | 9415 (39%) | 1625 (17%) | 982 (60%) | 643 (40%) | 672 (41%) | 440 (27%) | 513 (32%) |
| **SAT** | 8976 (37%) | 1373 (15%) | 814 (61%) | 525 (39%) | 756 (56%) | 471 (35%) | 112 (9%) |
| **BAT** | 8687 (36%) | 1409 (16%) | 665 (47%) | 744 (53%) | 494 (35%) | 613 (44%) | 302 (21%) |
| **Heart** | 7748 (32%) | 1233 (16%) | 646 (52%) | 587 (48%) | 434 (35%) | 397 (32%) | 402 (33%) |
| **Muscle** | 7145 (29%) | 1257 (18%) | 675 (54%) | 582 (46%) | 524 (42%) | 475 (38%) | 258 (21%) |
| **VAT** | 7019 (29%) | 981 (14%) | 525 (54%) | 456 (46%) | 434 (44%) | 356 (36%) | 191 (20%) |
| **Plasma** | 5469 (23%) | 1061 (19%) | 402 (38%) | 659 (62%) | 330 (31%) | 397 (37%) | 334 (32%) |

**VAT** (visceral adipose tissue), **SAT** (subcutaneous adipose tissue), **BAT** (brown adipose tissue)
<sup>#</sup>The percentage is based on the total number of molecular features detected in all the tissues from the ionization modes (*i.e.*, 24,294).
\*Defined as having a fold change $\geq 1.3$ , $p$ -value $\leq 0.05$ , and $q$ -value $\leq 0.05$ .
<sup>†</sup>The percentage is based on the tissue-specific number of the total significantly changed molecular features.

**Supplementary Table 2.** List of annotated compounds from volcano plots and k-mean cluster analyses (excel format).

